# Contrasting patterns of water use efficiency and annual radial growth among European beech forests along the Italian peninsula

**DOI:** 10.1101/2023.11.15.567154

**Authors:** Paulina F. Puchi, Daniela Dalmonech, Elia Vangi, Giovanna Battipaglia, Roberto Tognetti, Alessio Collalti

## Abstract

Tree mortality and forest dieback episodes are increasing due to drought and heat stress. Nevertheless, a comprehensive understanding of mechanisms enabling trees to withstand and survive droughts remains lacking. Our study investigated basal area increment (BAI), and δ^13^C- derived intrinsic water-use-efficiency (_i_WUE), to elucidate beech resilience across four healthy stands in Italy with varying climates and water availability. Additionally, fist-order autocorrelation (AR1) analysis was performed to detect early warning signals for potential tree dieback risks during extreme drought events.

Results reveal a negative link between BAI and vapour pressure deficit (VPD), especially in southern latitudes. After the 2003 drought, BAI decreased at the northern site, with an increase in δ^13^C and _i_WUE, indicating conservative water-use. Conversely, the southern sites showed increased BAI and _i_WUE, likely influenced by rising CO_2_ and improved water availability. In contrast, the central site sustained higher transpiration rates due to higher soil water holding capacity (SWHC). Despite varied responses, most sites exhibited reduced resilience to future extreme events, indicated by increased AR1.

Temperature significantly affected beech _i_WUE and BAI in northern Italy, while VPD strongly influenced the southern latitudes. The observed increase in BAI and _i_WUE in southern regions might be attributed to an acclimation response.

## Introduction

Forest ecosystems are facing significant challenges due to anthropogenic climate change^1^. The combination of reduced water availability and rising temperatures directly impacts the process of photosynthetic carbon assimilation, thereby reducing forest carbon sequestration. This could potentially lead to negative feedback on carbon balance^2^. Furthermore, hotter droughts have caused substantial alterations in forest structure and function, affecting tree growth performance and triggering episodes of dieback and tree mortality^3^. In addition, climatic models predict that the frequency, duration, and intensity of extreme droughts will increase in the future, so it is crucial to a better understanding of how forests are going to cope with these extreme climatic conditions^4^.

Despite the importance of identifying suitable tree species and future management practices in response to climate change, our understanding of species-specific physiological responses and site- and species-specific vulnerabilities to drought-induced tree mortality during extreme droughts remains incomplete^5^. This gap is especially critical for European beech (*Fagus sylvatica* L.), one of the most distributed, ecologically and economically significant tree species in Europe. This species comprises 17% of all broadleaf tree stands in Italy^6^ and is one the most affected by extreme events occurring during the initial vegetative phase across the Italian peninsula^7^.

Given the anticipated that climate change will exert a significant influence on both regional and local drought patterns in the Mediterranean region^8^. In particular, mountain-Mediterranean beech forests would face increased vulnerability due to their location in the southernmost distribution of this species’ range^9^. Consequently, predicting resilience and adaptation across its distribution has become a prioritized goal.

Recent studies have shown that prolonged heat and drought events can have detrimental effects on both hydraulic function and carbon use in trees^10^. Understanding these physiological mechanisms is crucial for comprehending how trees respond to drought, as they directly influence water use regulation. For instance, isohydric species adopt a conservative behaviour by closing stomata to minimize water loss, thereby reducing photosynthetic activity, and increasing the risk of carbon starvation^11^. On the other hand, anisohydric species adopt an opportunistic behaviour, exhibiting higher transpiration rates even when soil moisture is low, leading to an elevated risk of hydraulic failure^12^.

Currently, there is contrasting information regarding how European beech forests respond to heat and drought events. Most studies on young beech stands have suggested a conservative response during droughts^13^. However, in a few studies, adult trees have conversely displayed opportunistic behaviour^9^. Therefore, it is crucial to exploit better the plasticity of this species in the water use strategies to determine the trajectories of species distribution and its resilience to a warming and drier climate^14^

Long-term changes in intrinsic water use efficiency (_i_WUE), i.e. the cost of fixing carbon per unit of water loss^15^, can be assessed by measuring carbon isotope composition in tree rings (δ^13^C). Tree- ring δ^13^C is equivalent to the ratio between photosynthesis (*A*) and stomatal conductance (*g*_s_) and this can vary, since both affect the ratio between CO_2_ partial pressure in leaf intercellular space and in the atmosphere^16^. Variations in _i_WUE, within and across tree species, have revealed a continuous ecophysiological gradient of plant water-use strategies ranging from “profligate/opportunistic” (low _i_WUE) to those considered “conservative” (high _i_WUE)^17^. For instance, studies in tree rings have shown that the increase of _i_WUE did not enhance tree growth^18^, however, others showed the opposite effect or both^18–20^. These indicators of hydraulic strategies and carbon discrimination provide valuable insights into the long-term impacts of climate change on forest health and the risk of tree mortality^21^.

On the other hand, recent studies have provided evidence that one of the primary mortality risk indicators in forests is growth reduction also occurring many decades before visible symptoms of decline, such as leaf discolouration, increased defoliation, and branch dieback^22,23^. Similarly, another proxy indicator of loss of resilience and thus increasing tree mortality risk is the autocorrelation, better called ‘early warning signal’ (EWS), which has been proposed to detect a critical transition in long-term time series after a perturbation, causing a critical slowing down of the capacity of recovery^24,25^. EWS can be highlighted as increasing autocorrelation and variance in tree growth, indicating loss of resilience and stability^24,26^. These changes have been observed in conifers; however, angiosperms did not show changes in these indicators, and this could be due to their capacity to recover after a stress-induced growth decline^22,23^. These findings highlight the importance of early monitoring in understanding forest resilience and adaptation to climate change.

This study aimed to assess the forest vitality of beech in response to drought stress by examining historical and recent growth patterns across the Italian peninsula, with a particular emphasis on water use strategies (conservative vs. opportunistic) at long-time scales. Secondly, we tested early warning signals of potential tree dieback by analyzing autocorrelation and variability patterns, as indicators of stand resilience and stability to future extreme events.

We hypothesized that beech populations in the southernmost distribution exhibit conservative behaviour as an acclimation strategy. This behaviour is characterized by _i_WUE being more responsive to VPD than those in the northern regions, reflecting a reduction in stomatal conductance to maintain a minimum midday water potential, and also a decline in intercellular CO_2_ concentration, but a more slowly decrease in photosynthetic rate. Although a drought-driven decline in photosynthetic rate may also occur, non-stomatal limitation was expected in populations with more opportunistic behaviour. Additionally, we expected to find varying degrees of growth reduction as an early warning signal of tree mortality risk across different sites, with the strongest signals in response to severe drought events.

## Materials and Methods

### Study Sites and Climate

Analyses were conducted at four sites along a ∼900 km latitudinal transect in pure European beech forests across the Italian Peninsula (Fig. 1, Table 1). The sites were Trentino-Alto Adige (hereafter abbreviated as ‘TRE’), Lazio (hereafter abbreviated as ‘LAZ’), Campania (hereafter abbreviated as ‘CAM’) and Calabria (hereafter abbreviated as ‘CAL’). All the stands analyzed had not been managed since the last 20-30 years.

**Fig. 1.**
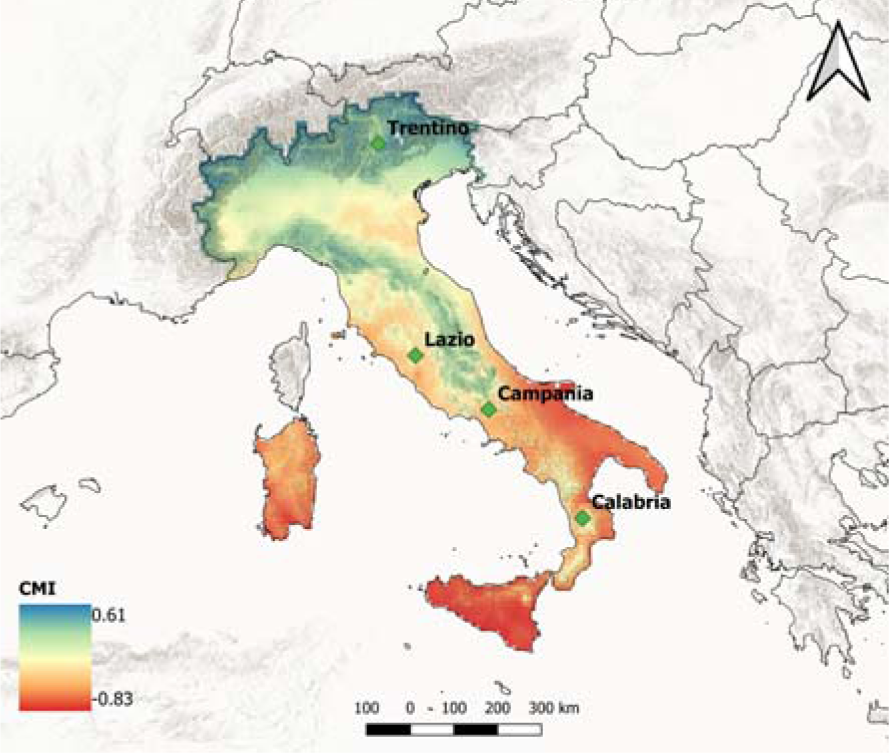
a) Map displaying the Climate Moisture Index (CMI = Precipitation / Potential EvapoTranspiration) across the Italian Peninsula, indicating humid and dry climate zones through positive (blue) and negative (red) CMI values, respectively. The index was calculated for the growing season (May-October) from 1965 to 2014. Green dots indicate the location of the four study sites where dendrochronological samples were extracted.

**Table 1.** Geographical and mean annual climate characteristics for the four sites.

| Site | Latitude<br>(N) | Longitude<br>(W) | Elevation<br>(m a.s.l.) | Mean<br>minimum<br>annual<br>temperature<br>(°C) | Mean<br>annual<br>temperature<br>(°C) | Mean<br>maximum<br>annual<br>temperature<br>(°C) | Annual<br>precipitation<br>(mm) |
| --- | --- | --- | --- | --- | --- | --- | --- |
| TRE | 46°12' | 11°16' | 1276 | 3.8 | 9.0 | 13.9 | 929 |
| LAZ | 42°24' | 12°12' | 1000 | 8.9 | 14.3 | 19.5 | 829 |
| CAM | 41°24' | 14°20' | 1140 | 6.3 | 9.8 | 13.1 | 825 |
| CAL | 39°19' | 16°23' | 1601 | 5.8 | 11 | 16 | 1003 |

The selection of the sites allowed the comparison of moisture availability across the Italian Peninsula (Fig. 1), using the Climate Moisture Index (CMI) calculation method explained in section 2.6. These sites differ along the latitudinal transect regarding both climatic conditions and soil types. From north to south, the mean annual temperature ranges from 9 to 14.1 °C, with the mean annual precipitation varying from 1003 to 825 mm, based on the E-OBS dataset (as explained in section E-OBS daily climate data, CMI and SPEI calculation; Supplementary Fig. S1)

The soil types from north to south are Andisols, Luvisols, and Inceptisols. Additionally, by examining soil texture data, we inferred variations in soil water holding capacity (SWHC) among these sites^27^ (Supplementary Table 1). Specifically, we inferred that the SWHC in TRE is relatively low, whereas LAZ exhibited a high SWHC. CAM also showed a high SWHC, while CAL presented a moderate SWHC^28^.

### Field sampling and processing dendrochronological data

During the period 2014-2018, a total of 174 beech trees were sampled at 1.3 m from the ground using a 5 mm increment borer (Supplementary Table 2). In the laboratory, wood cores were air/dried and polished with sandpaper of successively increasing grains to visualize the ring boundaries. Ring widths were measured to a precision of 0.01 mm using the TSAP measuring device (Rinntech). Tree-ring (TRW) series were then visually cross-dated using standard dendrochronological methods^29^ and checked for dating accuracy and measurement errors with the COFECHA program^30^.

Later, tree growth measurements were converted to basal area increment based (BAI) based on the distance between the outermost measured ring (pith) and the last ring of the tree (i.e., the ring next to the bark), using the following formula:

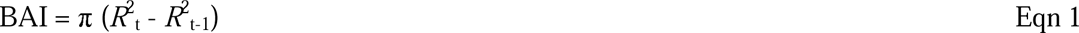

where *R*_t_ the tree’s radius at the end of the annual increment, and *R*_t-1_ is the tree’s radius at the beginning of the annual increment. This method assumes a circular cross-section, and the mean BAI of defined periods can be compared over time, as it is not affected by biological trends^31^ and it is more tightly related to stem biomass compared to TRW. We worked with mean non-standardized BAI values to preserve the long-term cumulative effects of climate on tree growth^19^. All analyses were restricted to the period covered by the youngest trees (at LAZ), i.e. from 1965 until 2014 (Supplementary Table 2). All computations were performed using the R-package ‘dplR’^32^.

### Water-use efficiency from carbon isotope discrimination

To compare long-term changes in _i_WUE among beech trees across the Italian Peninsula, we measured ^13^C/^12^C isotope ratios in the TRW. Ten samples per each stand presenting the best cross-dating (GLK > 0.70) with the corresponding average chronology, were selected for stable isotope analyses^33^ and they were annually dissected using a razor blade under a binocular microscope for the period 1965-2014.

Wood samples were milled to a fine powder (ZM 1000; Retsch), weighed 0.05-0.06 mg of wood for carbon isotope analyses and encapsulated in tin capsules.

The isotope composition was measured at the IRMS laboratory of the University of Campania “Luigi Vanvitelli” by using mass spectrometry with continuous flow isotope ratio (Delta V plus Thermo electron Corporation). The standard deviation for repeated analysis of an internal standard (commercial cellulose) was better than 0.1‰ for carbon. The δ^13^C series were corrected for the fossil fuel combustion effect for anthropogenic changes in the atmospheric δ^13^C composition (δ^13^C_atm_)^34^.

Isotopic discrimination between the carbon of atmospheric CO_2_ and wood carbon to determine _i_WUE can be calculated starting from the δ^13^C of the plant material (δ^13^C_tree_), which is related to atmospheric δ^13^C (δ^13^C_atm_) and the ratio c_i_/c_a_, according to Farquhar et al.^16^:

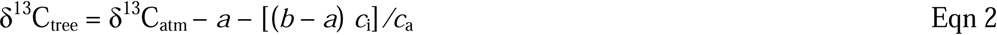

where *a* is the fractionation factor due to ^13^CO_2_ diffusion through stomata (4.4‰), and *b* is the fractionation factor due to Rubisco enzyme during the process of carboxylation (27‰)^34^. Therefore, we can calculate *c*_i_ by using the formula:

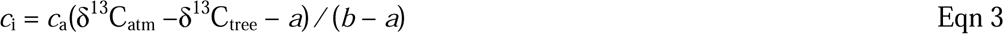

Finally, the _i_WUE can be calculated as follows:

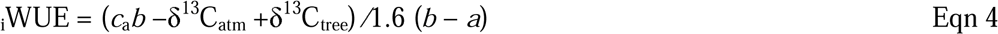

However, the _i_WUE should not be considered equivalent to instantaneous WUE at leaf level, which is the ratio of assimilation to stomatal conductance and considers the atmospheric water demand^15^. Thus, the equation used is the “simple” form of isotopic discrimination that does not include effects due to mesophyll conductance and photorespiration, which were unavailable for the study species.

We used δ^13^C_atm_ values from Belmecheri & Lavergne^35^. We obtained the atmospheric concentration of CO_2_ from the Mauna Loa station data (http://www.esrl.noaa.gov/).

### E-OBS daily climate data, CMI and SPEI calculation

Daily climate data for precipitation (P), minimum (T_min_), mean (T_mean_), and maximum (T_max_) temperature, as well as relative humidity (RH), were extracted from the E-OBS dataset on a regular 0.1-degree grid (Table 1). The data were obtained as netCDF files from (http://surfobs.climate.copernicus.eu/dataaccess/access_eobs.php). Using the RH and temperature data, the vapour pressure deficit (VPD) in hPa was calculated based on the Tetens formula^36^.

The Climate Moisture Index (CMI)^37^ represents the relationship between plant water demand and available precipitation. The CMI indicator ranges from –1 to +1, with wet climates showing positive CMI and dry climates negative CMI. CMI was calculated as follows:

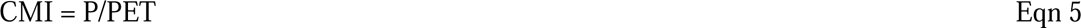

Where P is the precipitation, and PET is the potential evapotranspiration. Specifically, CMI = (P/PET) –1 when P<PET and CMI = 1–(PET/P) when P>PET, to recast the limit to –1<CMI<1.

PET can be calculated through the Hargreaves equation (Hargreaves, 1985), modified by Allen^38^:

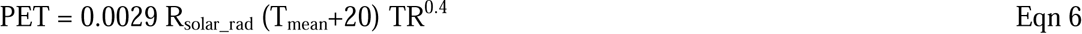

Where R_solar_rad_ is the extraterrestrial solar radiation, T_mean_ in Celsius degree and TR is the temperature range (T_max_–T_min_).

CMI was calculated for the growing season (May-October) using the E-OBS v. 27.0 (https://surfobs.climate.copernicus.eu/dataaccess/access_eobs.php#datafiles) daily products (T_min_, T_max_, precipitation, and global solar radiation) at 0.1 deg spatial resolution and averaged over the period 1965-2014. E-OBS global solar radiation at the surface was converted to extra-terrestrial solar radiation with the ‘envirem’ R-package^39^.

Additionally, to quantify drought severity, we calculated the Standardized Precipitation-Evapotranspiration Index (SPEI), based on a statistical transformation of the climatic water balance, i.e. precipitation *minus* potential evapotranspiration (P-PET). The multiscalar drought index was calculated at different time scales (from 1 to 24 months, Supplementary Fig. 2) for the period 1965- 2014 (constrained to the youngest site LAZ) in R using the ‘SPEI’ package^40^

Later, to assess the relationships between climate and BAI and stable isotope for the period 1965– 2014, we calculated Pearson’s correlations between monthly P, T_mean_, VPD, and SPEI (1-3-6-9-12- 18 and 24 months) series from previous _(t-1)_ and current year _(t)_, using monthly response function in the ‘DendroTools’ R-package^41^.

### Growth trends and climate response

We used Generalized Additive Mixed Models (GAMM) to study the long-term annual BAI and their responses to changing climatic conditions, particularly concerning water balance within the growing season (May-October) using SPEI indexes at the four study sites. We tested SPEI drought index at 1,3,6,9,12,18, and 24 months as the potential influence of drought on BAI. GAMM is a flexible semiparametric method that allows the simultaneous modelling of linear and nonlinear relationships between the response variable as a function of some explanatory variables that allows the treatment of autocorrelation and repeated measures^42^. The variables included in the model were the following:

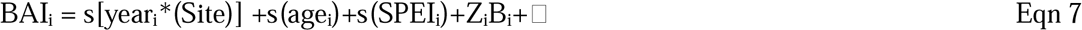

Where the BAI*_i_* of a tree_i_ were modelled as a function of calendar year, individual tree age and SPEI per site. In addition, given that BAI represents multiple measurements performed in each tree, tree identity (*Z_i_B*_i_) is regarded as a random effect (*Z_i_*and *B_i_* indicate matrix variables and related coefficients). Thin plate regression splines (*s*) were used to represent all the smooth terms, with a degree of smoothing determined by internal cross-validation^42^. We ranked all the potential models that could be generated using different explanatory variables and different levels of smoothing according to the Akaike Information Criterion (AIC). Finally, we chose the model with the lowest AIC^43^. The time scale that best explained the variability in BAI was the 18-month SPEI (for the growing season May-October). The GAMMs were performed and fitted using the function ‘gamm’ in the ‘mgcv’ R-package^42^.

### Early warning signals of forest dieback

For assessing stand resilience for each site and each time series of BAI, we computed the first-order autocorrelation (AR1) and the standard deviation (SD) over the period 1965 to 2014 using a 15-year moving window (30% of the entire time series). These metrics are widely recognized indicators of changes in time series and proximity to critical transitions to new states^22,24,25^. The trend of AR1 and SD metrics over the considered temporal window was computed by means of the non-parametric Mann-Kendall Tau statistics. For each site, the significance of a positive (or negative) AR1 and SD trend was tested with a one-sided t-test. We employed the R-package ‘early warnings’^26^ to compute the selected metrics.

## Results

### Climate trends and drought variability

Annual precipitation (P) has increased significantly at TRE and CAM sites (*P* < 0.01, Supplementary Fig. 2a), while LAZ showed a reduction in P trend during the period from 1965 to 2014 (*P* < 0.05, Supplementary Fig. 2a), and CAL did not present any trend in P pattern. Notably, T_min_ increased significantly in TRE, LAZ, and CAM (*P* < 0.001, Supplementary Fig. 2b), whereas in the southern site (CAL), T_min_ presented a pronounced decrease (*P* < 0.01, Supplementary Fig. 2b). Simultaneously, both T_mean_ and T_max_ exhibited a substantial and significant increase across all sites (*P* < 0.01, Supplementary Fig. 2c,d). Interestingly, only at the northernmost site (TRE), VPD increased drastically and significantly during the 2000s (*P* < 0.001, Supplementary Fig. 2e), while at the southernmost site (CAL) VPD showed the opposite pattern (Supplementary Fig. 2e, P < 0.001).

As for the P trend, the SPEI index showed an increase in water availability in recent years across sites, although not significant, except for LAZ, which showed a negative trend (*P* < 0.05, Fig. 2, Supplementary Fig. 3). Notably, the SPEI-derived drought index showed the widespread impact of the 2003 drought across all sites, more evident at CAM (Fig. 2, Supplementary Fig. 3).

**Fig. 2.**
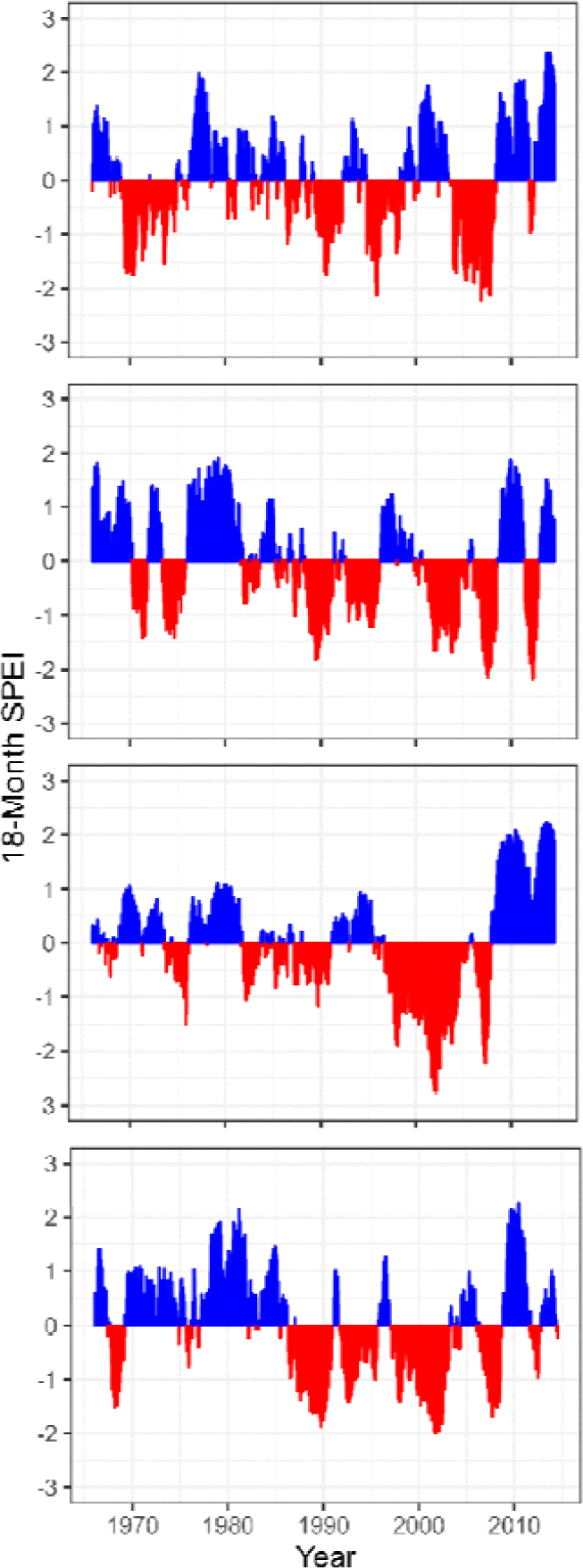
Standardized 18-Month SPEI at the four study sites (TRE, LAZ, CAM, and CAL) for the 1965–2014 period. Negative (red) and positive (blue) values indicate drier and wetter conditions, respectively.

### *Long*-term growth patterns of European beech across the Italian Peninsula

Mean TRW, the highest and the lowest growth rates were observed in LAZ and CAM sites, respectively with statistically significant differences. Conversely, TRE and CAL showed similar growth rates values (*P* < 0.05, Supplementary Table 3). The age distribution of tree populations exhibited notable differences across the four sites, with LAZ featuring the youngest trees and CAM the oldest trees (*P* < 0.05, Supplementary Table 3).

BAI exhibited a significant decline, particularly pronounced in the relatively northern sites (TRE and LAZ), following the drought of 2003 (Fig. 3). In contrast, CAM presented a steady increase in BAI, while in CAL, BAI decreased after 2010 (Fig. 3).

**Fig. 3.**
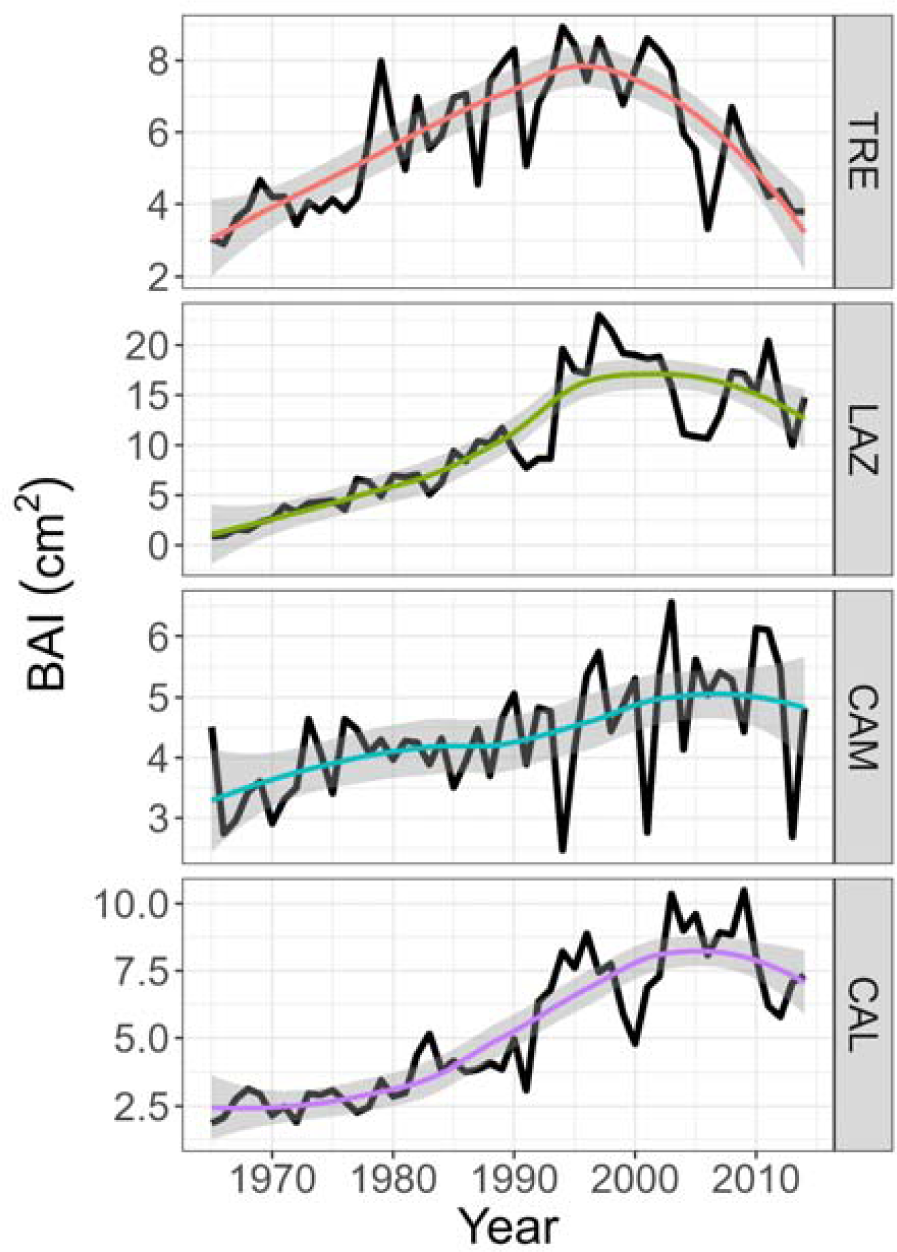
Long-term growth of basal area increment (BAI) of the four sites along a latitudinal gradient (north to south) for the period 1965–2014. Colour lines for each site indicate the linear model, shaded areas represent 95 % confidence intervals.

### Growth response to climate variables

Basal area increment exhibited significant relationships with climatic variables in all study sites (Fig. 4). Overall, BAI was positively correlated with monthly P and T_mean_ and, notably, strongly negative correlations with VPD were evident from May to September. This negative VPD correlation intensified toward the southern sites (Fig. 4).

**Fig. 4.**
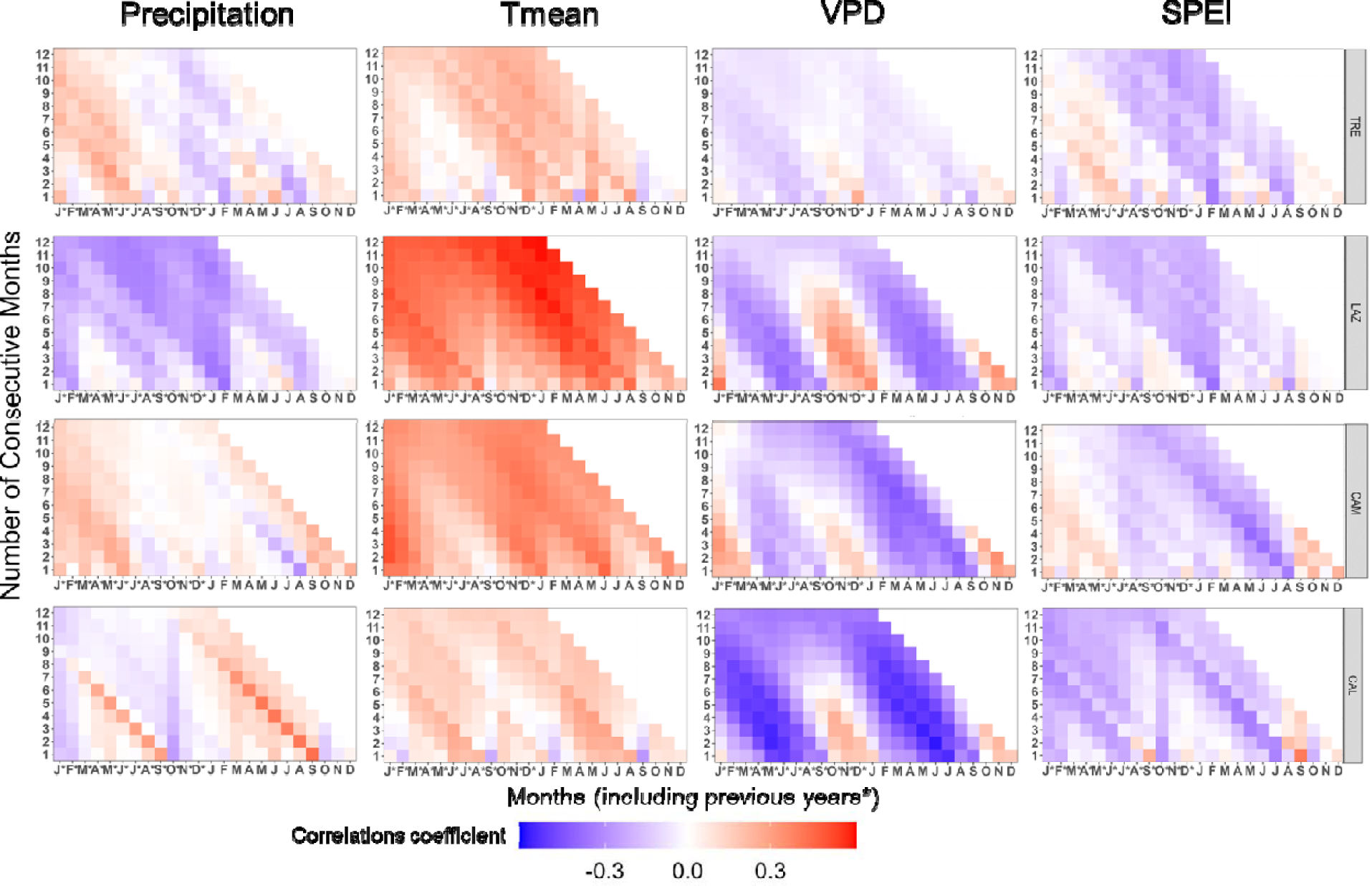
Pearson’s running correlations between BAI with monthly precipitation, mean temperature, VPD, and SPEI1 for the current and the previous year (*) over the period 1965-2014 at each site. The *y-axis* represents the time window in months. Colours (see the key) represent correlation coefficients that are significant at the level of *r* = 0.279 (*P* < 0.05).

At TRE, BAI was positively correlated with monthly P from May to July, with a stronger effect when considering P values in the previous year. Additionally, BAI correlated positively with May T_mean_ of the current year, instead of showing weak negative correlations with VPD and SPEI (Fig. 4).

LAZ showed a strong positive correlation between BAI and T_mean_ from March to November (previous and current year). Conversely, strong negative correlations with VPD from May to September and weak negative correlations with P and SPEI were found in August (Fig. 4).

AT CAM, a positive response of BAI to P of July of the previous years, and a strong positive correlation with T_mean_ during May to July, were observed, while a strong negative response to VPD from March to September (more evident in the current year) was found.

At CAL, BAI showed strong positive correlations with October P and with T_mean_ from March to August. In contrast, BAI displayed a strong negative correlation with VPD from March to September (current and previous year). Similarly, negative scattered correlations with SPEI were observed during summer at the southern sites (CAM and CAL, Fig. 4).

### Growth trends of beech

The GAMMs revealed different BAI trends of beech among the four sites (Fig. 5, Supplementary Table 4). GAMM showed a monotonic increasing growth trend among the sites; however, they started to diverge in the mid-1990s. Notably, the northernmost site (TRE) started to decline earlier than the other sites (Fig. 5). Secondly, LAZ exhibited the highest increase, followed by a drastic decline during the 2000s – a similar pattern was also observed in CAL. In contrast, at CAM trees demonstrated a steady increase in BAI over the observed period (Fig. 5).

**Fig. 5.**
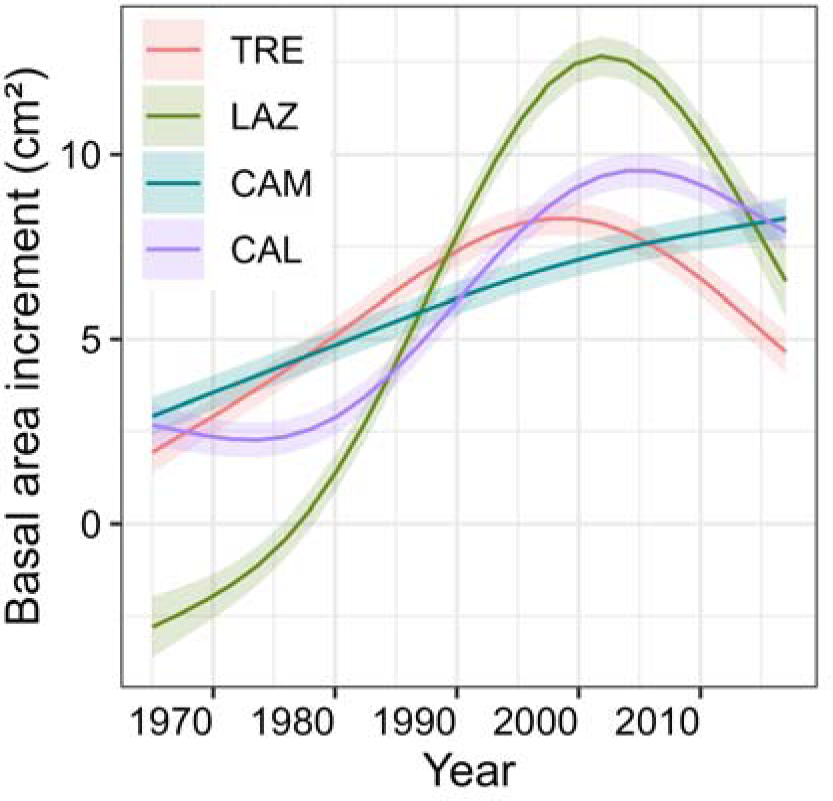
Growth trends of basal area increment trends of beech for the four sites. Trends were based on the best-fitted generalized additive mixed models (GAMM) for the period 1965-2014.

### Long-term carbon isotope chronologies and water use strategies

At the southernmost site (CAL), trees presented the highest increase of δ^13^C values that translate in an increase of _i_WUE (Table 3, Fig. 6). On the contrary, CAM showed the lowest value of _i_WUE (Table 3, Fig. 6). LAZ and TRE on average presented similar δ^13^C and _i_WUE (Table 3).

**Fig. 6.**
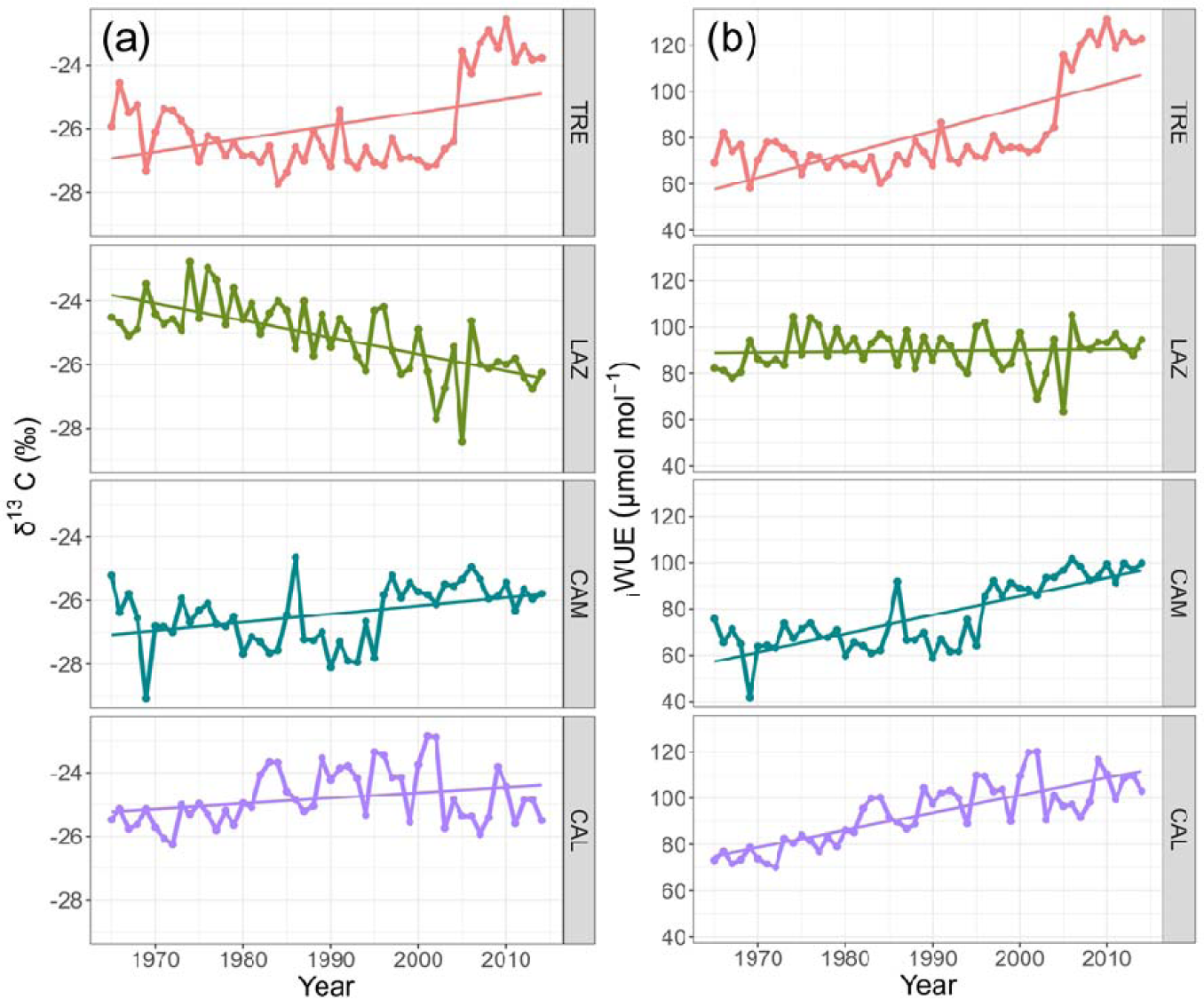
Trends of: a) δ^13^C (‰), and b) _i_WUE, and fitted linear trends for the period 1965–2014 in four stands across a latitudinal gradient in Italy.

**Table 3.**
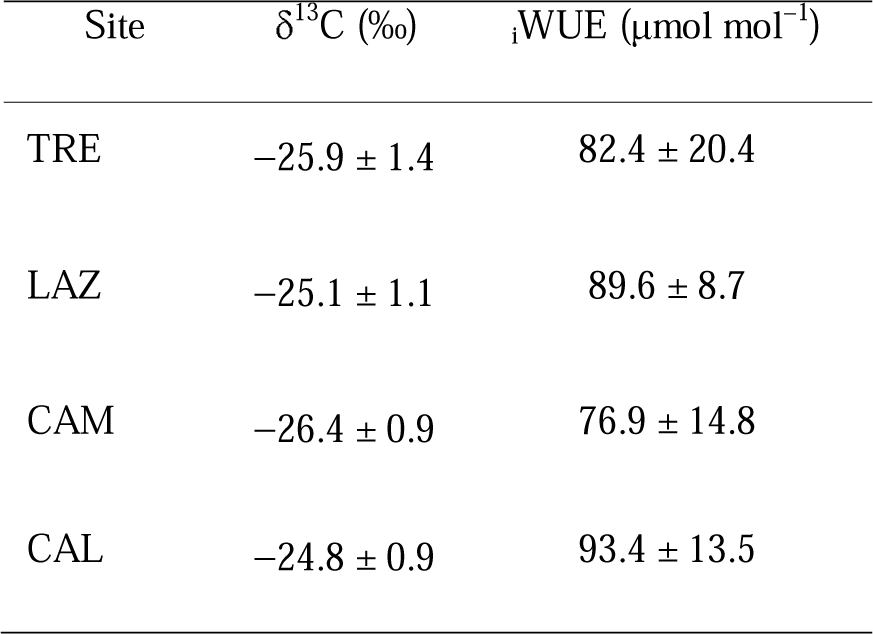
Statistics of mean δ^13^C and _i_WUE of the tree-ring width series of beech per site for the period 1965–2014. Data are mean values ± SE.

For most sites, δ^13^C showed a positive and significant trend over time (*P* < 0.05, Fig. 6), except for LAZ, which showed an opposite pattern during the period 1965–2014 (*P* < 0.001, Fig. 6). In the northernmost site (TRE), the δ^13^C and _i_WUE, started to increase sharply after the drought of 2003.

Similarly, the southern sites CAM and CAL presented a steady increase in _i_WUE (*P* < 0.001, Fig. 6). On the contrary, LAZ did not present any significant trend (*P* = 0.701).

In the southern sites, CAM and CAL, we observed significant positive relationships between _i_WUE and BAI (*P* < 0.001, Fig. 7). On the contrary, at the northern site (TRE), we observed the opposite trend pattern; however, this trend was not significant (*P* > 0.05). AT LAZ, no relationship was found between _i_WUE and BAI (Fig. 7).

**Fig. 7.**
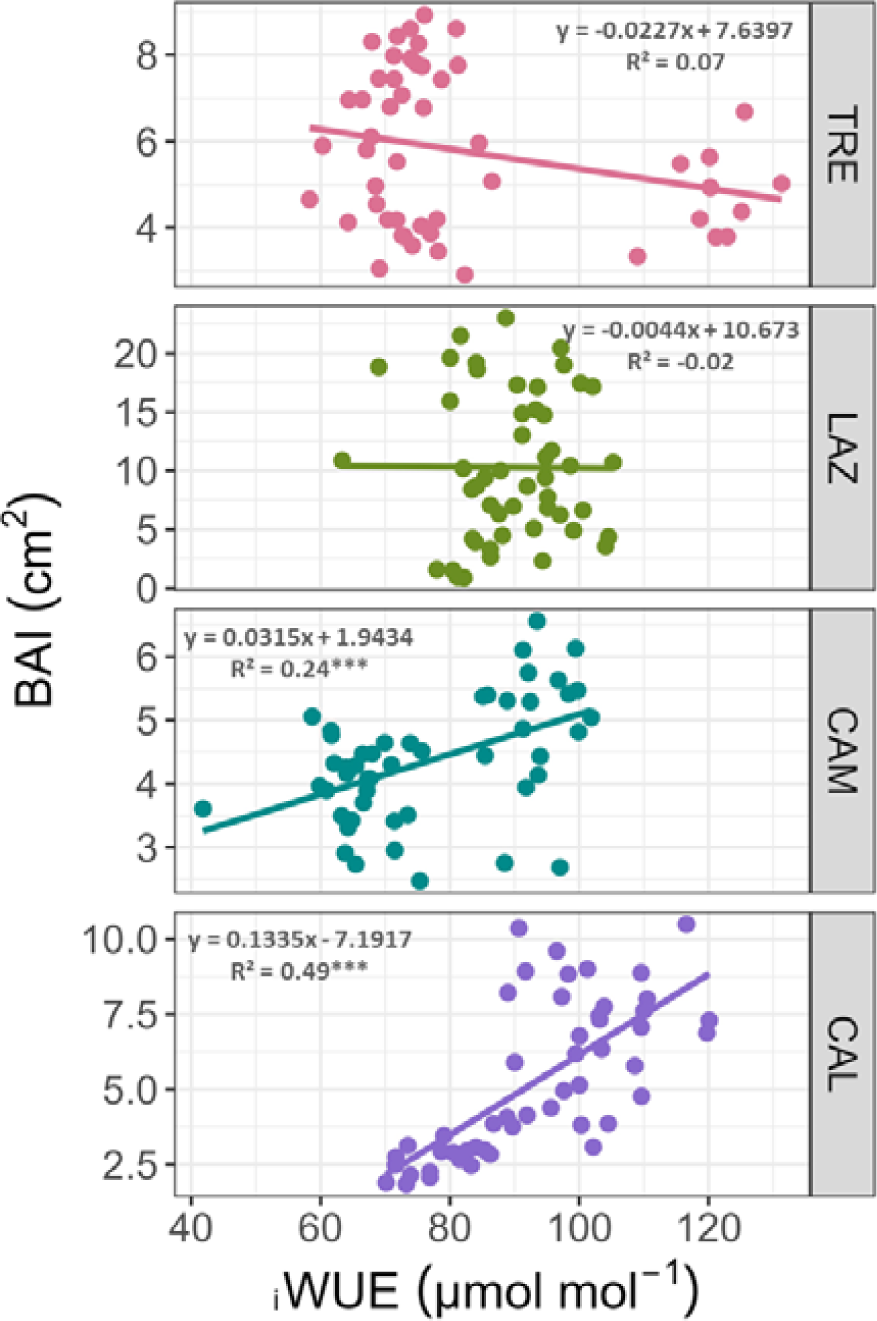
Relationship between annual BAI and _i_WUE in beech across the Italian Peninsula for the period 1965-2014. Linear regressions and the equations are indicated for each site. Significance values are encoded by \*\*\**P* < 0.001.

### ***δ****^13^C*, _i_WUE, and climate relationship

Carbon isotope composition (δ^13^C) and _i_WUE showed a similar relationship with climate variables. However, _i_WUE presented stronger correlation with climate than δ^13^C (Fig. 8, Supplementary Fig. 4). An exception was observed at LAZ, where δ^13^C showed a negative and significant correlation with T_mean_ compared to _i_WUE (Fig. 8, Supplementary Fig. 4).

**Fig. 8.**
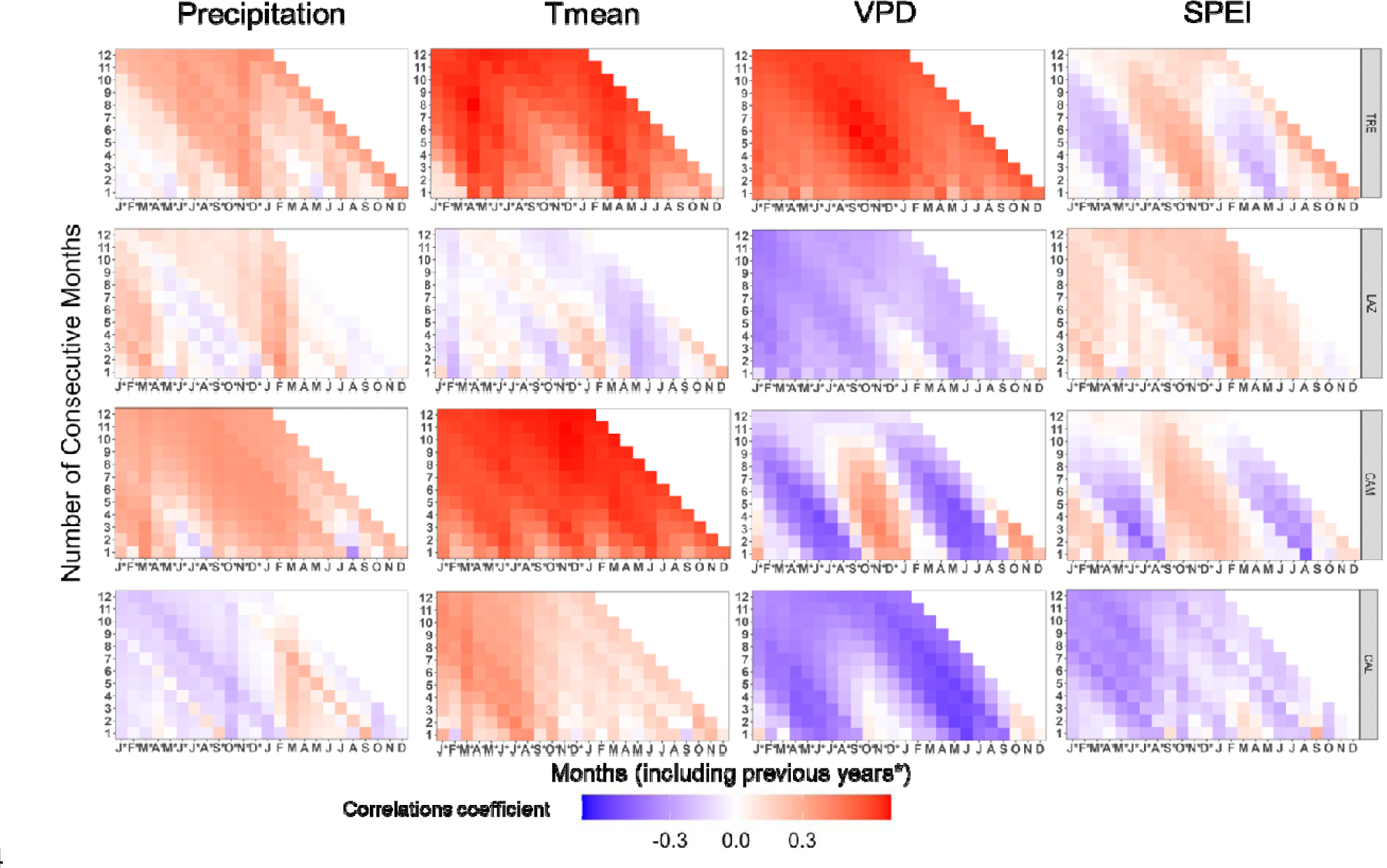
Pearson’s running correlations between _i_WUE with monthly precipitation, mean temperature, VPD, and SPEI1 for the current and the previous year (*), over the period 1965-2014 at each site. The *y-axis* represents the time window in months. Colours (see the key) represent correlation coefficients that are significant at the level of *r* = 0.279 (*P* < 0.05)

At the northernmost site, _i_WUE showed significant and positive correlations with T_mean_ and VPD of the previous and current year, while negative and scattered correlations with P and SPEI of April and May were observed (Fig. 8).

At LAZ, _i_WUE was negatively and significantly correlated with VPD from March to November of the current and previous year.

At CAM, _i_WUE exhibited a positive and significant correlation with P from March to May (previous and current year), while T_mean_ of the current and previous year was positively and significantly correlated with _i_WUE. On the contrary, _i_WUE correlated strongly and negatively with VPD from May to August (current and previous year) and with SPEI in August (Fig. 8).

At the southernmost site, _i_WUE correlated positively with P from March to September (current year), and strongly and positively with T_mean_ from March to August of the previous year. At CAL, VPD and SPEI exhibited strong and negative correlations with _i_WUE from May to June (current and previous year, Fig. 8).

### Early warning signals of declining forest resilience

The statistical analysis of the BAI time series performed to detect EWS on beech forests revealed contrasting results among the sites (Fig. 9a,b). In TRE and LAZ, BAI showed a rise in AR1 among trees, which started to increase after the 2003 drought in TRE, while in LAZ already during the 1990s (Fig. 9a). In contrast, CAM showed a significant steady decrease in AR1. No significant autocorrelation trend was found at CAL, Nevertheless, the standard deviation (SD) started to rise by the end of the 1980s (Fig. 9b). A significant increase in SD of the BAI signal was observed across all the sites.

**Fig. 9.**
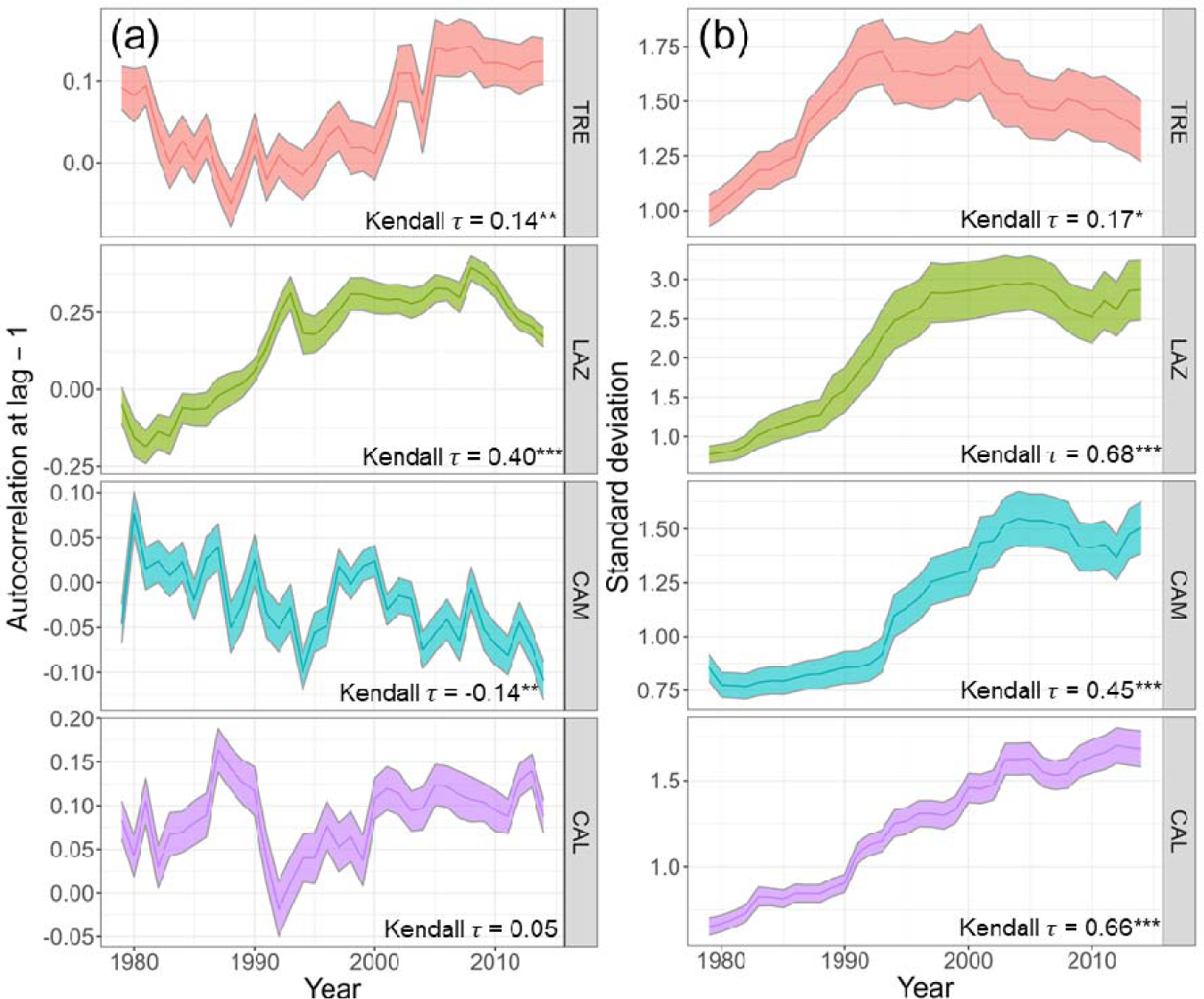
Early warning signals: a) AR1, first-order autocorrelation, b) SD, obtained using a 15-year moving window for basal area increment (BAI) of *Fagus sylvatica* for four study sites for the period 1965-2014. The statistics of BAI were calculated using the residuals of the time series after removing the low-frequency signal (Gaussian filter) using 15-year-long windows (e.g. 1979 corresponds to the interval 1965-2014). The Kendall statistics indicate the strength of trends along the time series for each variable and site. For each site, the bold line represents the mean of the statistics among trees, and the shaded area is the standard error. (* *P* < 0.05, ** *P* < 0.001, *** *P* < 0.001).

## Discussion

Our analysis revealed contrasting long-term growth responses of European beech across the Italian Peninsula, closely associated with local climate and site conditions. The northern sites (TRE and LAZ) showed a decrease in BAI trends after the severe drought event in 2003, while the southernmost site (CAL) exhibited a growth decline after 2010. In contrast, CAM displayed a steady increase in growth over the 50 years analysed, likely due to increased precipitation in the last decades.

Our findings confirmed that European beech in the northern sites might be more susceptible to die-off, even without visible decline symptoms (such as branch dieback or decolouration of leaves). Trees exhibited greater growth sensitivity to VPD during summer, and this effect became more pronounced at the southernmost site. VPD can be used to estimate atmospheric water status and is one of the most important environmental factors influencing plant growth^44^. Elevated VPD, associated with dry conditions, impacts stomatal conductance and the balance between carbon assimilation and water loss^10,45^. This indicates that drought, driven by enhanced evapotranspiration, will play a critical role throughout the beech forest’s latitudinal range in Italy.

Recent global-scale research by Yuan et al.^46^ highlighted the increase of VPD as a major atmospheric driver affecting forest productivity by imposing water stress on photosynthesis. Water-use strategies, particularly conservative/opportunistic responses within and across species, have been closely linked to soil moisture availability ^17^. Higher VPD and temperature accelerate soil moisture depletion causing a significant reduction in carbon uptake^47^, elevating the risk of drought-induced dieback through hydraulic failure and/or carbon starvation^8,48^.

Our results demonstrate a significant increase in VPD after the 2003 drought event in TRE, however we found a weak negative correlation between VPD and BAI. We can hypothesise this might be attributed to a lower soil water holding capacity (SWHC) at this site, potentially increasing vulnerability to growth decline, as observed in our GAMM model. Conversely, LAZ exhibited higher SWHC, likely contributing to higher transpiration rates and growth compared to other sites. CAM and CAL sites presented moderate SWHC and a declining VPD trend, indicating less stress than the TRE site. The GAMM model integrated responses to the SPEI index and individual age of each tree, thus, we speculate that hydraulic strategies under drought significantly impact long-term growth rates, reflecting site-specific and ontogenic plasticity responses of the species. These findings may suggest that young beech trees initially benefit from favourable climate conditions and higher transpiration rates; however, this advantage depends on soil water availability and makes them susceptible to rapid declines in growth during extreme drought events, as already observed in Switzerland^49^.

Our study, to our knowledge, is the first to show evidence of the negative impact of VPD on basal area increment in beech forests across the Italian peninsula. This correlation was evident in all sites but was even stronger at southern latitudes. In contrast, previous studies in mature beech stands did not find a significant climate correlation, attributed to the species’ mast-seeding behaviour and sensitivity to late frosts at the beginning of the growing season^50^. Other studies have identified lagged climate correlation with masting^51^. Additionally, Zimmermann et al.^52^, in central Germany, found that beech growth was highly sensitive to summer temperatures and extreme drought events after the 1980s.

European beech has commonly been classified as an opportunistic species, capable of maintaining higher transpiration rates even in relatively dry soil conditions^9^; however, this strategy increases the risk of cavitation^10^.

Our findings indicated that temperature and VPD emerged as primary drivers of _i_WUE in TRE, while VPD played a dominant role in the southern sites. However, in LAZ, _i_WUE did not exhibit a clear correlation with climate variables. This complex relationship highlights the interaction between VPD, stomatal conductance, and photosynthesis, as high VPD initially reduces stomatal conductance but not net CO_2_ assimilation rate, resulting in increased _i_WUE. Nevertheless, severe VPD-induced stomatal conductance restrictions, combined with declining soil moisture and other non-stomatal limitations, ultimately reduce photosynthetic rate and may lead to declining _i_WUE as VPD continues to rise^53^. Thus, the overall relationship between _i_WUE and VPD is likely hyperbolic^54^, and the sensitivity of photosynthesis to VPD will likely be weaker than the sensitivity of conductance to VPD.

Our study highlights contrasting _i_WUE of beech across the Italian peninsula. We observed an increase of δ^13^C and _i_WUE values in TRE, CAM, and CAL, indicating a conservative water use strategy when water availability is low. In contrast, LAZ exhibited a decrease of δ^13^C, suggesting an opportunistic response with stable _i_WUE regardless of the moisture condition. While at LAZ, changes in photosynthetic rate and stomatal conductance appeared to occur in the same direction with similar magnitude, at TRE, CAM, and CAL, stomatal conductance appeared to decrease proportionally more than photosynthetic rate, or the latter remaining stable or increasing with declining stomatal conductance. Thus, our findings confirm that water use strategies employed by beech are mostly site-specific and influenced by microclimatic conditions and soil water availability^34^, consistent with our hypothesis and consistent with prior research^18^.

Interestingly, our results indicate that higher mean _i_WUE did not result in an increase in the basal area increment on beech^55–57;^ instead, we observed contrasting responses consistent with previous studies^18,19^. Notably, the northern site displayed a drastic increase in _i_WUE after the 2003 drought event, coinciding with elevated VPD and temperature that may have led to stomatal closure (*g*_s_) and reduced photosynthesis (*A*), suggesting that the growth decline in this site was might triggered by intensified evapotranspiration and the lower SWHC as observed in other sites by others (e.g.^56,58^). At LAZ, there was no relationship between _i_WUE and growth, which can be explained by higher SWHC allowing higher transpiration rates and metabolic respiration, resulting in greater losses of photosynthetic assimilates, especially at higher temperatures^55,57^. Interestingly, in the southern sites, the increase of _i_WUE enhanced growth. This discrepancy may be attributed to the adaptation of beech trees in the southernmost distribution to water stress and high VPD^59,60^, suggesting that high _i_WUE is an adaptative trait^61^. Consequently, we can infer that the observed “conservative strategy”- characterized by low stomatal conductance and constant CO_2_ assimilation rate that enhanced growth – at CAM might be explained by a positive CO_2_ fertilization effect or long-term acclimation to elevated CO_2_^20^. Similar findings were reported in mature European beech stands in Spain, where an increased sensitivity to drought was observed across the southern range-edge distribution^18^. Recently, Qi et al.^62^ in China revealed varying water use strategies among larch trees. Mature trees presented a more ‘conservative strategy’ (low *g*_s_, constant assimilation rate (*A*)), whereas young trees maintained constant *g*_s_ and high *A*, indicating an opportunistic behaviour. Similar to our study, mature trees displayed greater sensitivity to atmospheric CO_2_ concentrations than their young counterparts.

It should be pointed out that a major influence of photosynthetic rate on intercellular CO_2_ concentration and δ^13^C, and the minor contribution of the regulation of stomatal conductance to _i_WUE, were observed in other studies on the same species^19^. These findings suggest unclear patterns of potential increased drought-related tree decline signs in mountain beech forests along the Italian latitudinal transect. Differences between leaf-level physiology and tree-ring level processes may arise, reflecting potential variations in the (re)translocation patterns of non-structural carbohydrates to organs^63,64^. Such complexities make tree-ring analysis a challenging tool for decipher tree responses to fluctuating seasonal conditions in the short term.

Our second hypothesis, linking the degree of growth reduction and tree growth instability to drought severity, was only partially confirmed by our findings. We observed an increase in the autocorrelation of the BAI signal across almost all sites, indicating an enhanced intrinsic biological memory within the trees and a loss of ecological system resilience^65,66^. Such increases have been linked to instabilities preceding external disturbances in various biological systems^25,26^, potentially leading to a transition to a new system state^67^. Recent studies investigating ecosystem productivity’s autocorrelation have identified reduced resilience in diverse forest types due to increased water limitations and climate variability^25,68^. Notably, after a severe drought, declining trees exhibited increases in BAI autocorrelation and variability before mortality^22^.

In line with our expectations, the northern site showed a significant increase in AR1 and a decline in BAI after the 2003 drought event. Conversely, CAM showed a decrease in AR1, suggesting greater resilience to changing climate conditions, despite experiencing the severe drought period of 2003. This higher resilience at CAM might be linked to the legacy of past conditions with less intra-annual variability in water availability compared to the TRE site, as supported by the SPEI multiscalar index (Fig. 2). TRE experienced several prolonged dry periods (i.e. SPEI<-1.5) before the 2000s. Additionally, relatively mature trees at CAM site might contribute to the population’s apparent stability^69^. Our data also revealed an increase in BAI series SD across all stands. While this variability encompasses both tree physiological signals and climate-driven vegetation dynamics, the co-occurrence rises in AR1, decline in BAI, and increase in SD in TRE, LAZ and CAL sites, may indicate a loss of system stability^24^. These results indicate potential challenges for trees to mitigate the impact of extreme events in the future.

Several studies have demonstrated that long-term rises in instability and reduced growth predispose European beech to elevated mortality risks under future climate-induced stress conditions^70,71^. The loss of stability in our study sites emphasizes the need for continuous monitoring and proactive management of beech forests, particularly in regions where climate change is projected to increase the frequency and severity of droughts. Ongoing monitoring enables early detection of tree mortality risks, facilitating timely interventions to protect and sustain these vital ecosystems with wide ecological amplitude.

## Conclusions

In this study, our goal was to advance the early prediction of mortality risk in healthy beech stands without, apparently, visible declining symptoms across the Italian Peninsula, analyzing growth and _i_WUE patterns, although the available evidence is not yet conclusive.

These findings highlight the importance of considering the plasticity and site-specific _i_WUE responses to varying environmental conditions and the impact of VPD on stomatal conductance when predicting the future of beech forests in the context of climate change. It is important to note that not all beech populations considered in this study exhibited an increase in _i_WUE in response to rising VPD. This variability reflects differing sensitivities to changes in environmental drivers and the plasticity of conservative to opportunistic water-use strategies.

Furthermore, our analysis of EWS reveals the loss of resilience after an extreme event, as notably observed at the TRE site. In the context of climate model projection, the increase in the frequency and severity of droughts, the ability to detect earlier tipping points of critical slowdowns in declining systems, and the potential for recovery to the current state or an alternative state remains uncertain.

Nonetheless, this research raises further questions, such as how to generalize the relationships between increased _i_WUE and conservative behaviour, thus explaining contradictory results obtained in tree ring studies on beech populations and assessing temporal changes in this functional trait. Further research considering young and mature trees and their physiological mechanisms, micro- site conditions, and genetics will also elucidate intraspecific variations in drought response. This knowledge is essential for developing effective conservation and future forest management strategies to ensure these crucial ecological and socio-economical ecosystems’ long-term health, vitality, and resilience.

## References

1. McDowell, N. G. et al. Pervasive shifts in forest dynamics in a changing world. Science 368, 1–10 (2020).

2. Pan, Y. et al. A Large and Persistent Carbon Sink in the World’s Forests. Science 333, 988–993 (2011).

3. Puchi, P. F., Camarero, J. J., Battipaglia, G. & Carrer, M. Retrospective analysis of wood anatomical traits and treeDring isotopes suggests siteDspecific mechanisms triggering araucaria araucana droughtDinduced dieback. Glob Chang Biol 27, 6394–6408 (2021).

4. Brodribb, T. J., Powers, J., Cochard, H. & Choat, B. Hanging by a thread? Forests and drought. Science (1979) 368, 261–266 (2020).

5. Allen, C. D., Breshears, D. D. & McDowell, N. G. On underestimation of global vulnerability to tree mortality and forest die-off from hotter drought in the Anthropocene. Ecosphere 6, 1–55 (2015).

6. Gasparini, P., Di Cosmo, L., Floris, A. & De Laurentis Editors, D. *Italian National Forest Inventory— Methods and Results of the Third Survey*. https://link.springer.com/bookseries/15088 (2022).

7. Martinez del Castillo, E., et al. Climate-change-driven growth decline of European beech forests. Commun Biol 5, (2022).

8. Adams, H. D., et al. A multi-species synthesis of physiological mechanisms in drought-induced tree mortality. Nat Ecol Evol 1, (2017).

9. Leuschner, C. Drought response of European beech (Fagus sylvatica L.)—A review. Perspect Plant Ecol Evol Syst 47, (2020).

10. Mcdowell, N. et al. Mechanisms of Plant Survival and Mortality during Drought: Why Do Some Plants Survive while Others Succumb to Drought? New Phytologist 178, 719–739 (2008).

11. Timofeeva, G. et al. Long-term effects of drought on tree-ring growth and carbon isotope variability in Scots pine in a dry environment. Tree Physiol 37, 1028–1041 (2017).

12. Petrucco, L., Nardini, A., Von Arx, G., Saurer, M. & Cherubini, P. Isotope signals and anatomical features in tree rings suggest a role for hydraulic strategies in diffuse drought-induced die-back of Pinus nigra. Tree Physiol 37, 523–535 (2017).

13. Walthert, L. et al. From the comfort zone to crown dieback: Sequence of physiological stress thresholds in mature European beech trees across progressive drought. Science of the Total Environment 753, (2021).

14. Gessler, A., Bottero, A., Marshall, J. & Arend, M. The way back: recovery of trees from drought and its implication for acclimation. New Phytologist 228, 1704–1709 (2020).

15. Seibt, U., Rajabi, A., Griffiths, H. & Berry, J. A. Carbon isotopes and water use efficiency: Sense and sensitivity. Oecologia 155, 441–454 (2008).

16. Farquhar, G. D., O’Leary, M. H. & Berry, J. A. On the Relationship between Carbon Isotope Discrimination and the Intercellular Carbon Dioxide Concentration in Leaves. Aust. J. Plant Physiol 9, 121–158 (1982).

17. Moreno-Gutiérrez, C., Dawson, T. E., Nicolás, E. & Querejeta, J. I. Isotopes reveal contrasting water use strategies among coexisting plant species in a mediterranean ecosystem. New Phytologist 196, 489–496 (2012).

18. Peñuelas, J., Hunt, J. M., Ogaya, R. & Jump, A. S. Twentieth century changes of tree-ring δ13C at the southern range-edge of Fagus sylvatica: Increasing water-use efficiency does not avoid the growth decline induced by warming at low altitudes. Glob Chang Biol 14, 1076–1088 (2008).

19. Tognetti, R., Lombardi, F., Lasserre, B., Cherubini, P. & Marchetti, M. Tree-ring stable isotopes reveal twentieth-century increases in water-use efficiency of fagus sylvatica and nothofagus spp. in Italian and Chilean Mountains. PLoS One 9, (2014).

20. Walker, A. P. et al. Integrating the evidence for a terrestrial carbon sink caused by increasing atmospheric CO2. New Phytologist 229, 2413–2445 (2021).

21. Gessler, A. et al. Drought induced tree mortality – a tree-ring isotope based conceptual model to assess mechanisms and predispositions. New Phytologist 219, 485–490 (2018).

22. Camarero, J. J., Gazol, A., Sangüesa-Barreda, G., Oliva, J. & Vicente-Serrano, S. M. To die or not to die: Early warnings of tree dieback in response to a severe drought. Journal of Ecology 103, 44–57 (2015).

23. Cailleret, M. et al. Early-warning signals of individual tree mortality based on annual radial growth. Front Plant Sci 9, 1–14 (2019).

24. Dakos, V. et al. Methods for detecting early warnings of critical transitions in time series illustrated using simulated ecological data. PLoS One 7, (2012).

25. Forzieri, G., Dakos, V., McDowell, N. G., Ramdane, A. & Cescatti, A. Emerging signals of declining forest resilience under climate change. Nature 608, 534–539 (2022).

26. Dakos, V., Van Nes, E. H., D’Odorico, P. & Scheffer, M. Robustness of variance and autocorrelation as indicators of critical slowing down. Ecology 93, 264–271 (2012).

27. Tomislav Hengl & Travis Nauman. Predicted USDA soil great groups at 250 m (probabilities) (Version v01) [Data set]. Zenodo. Predicted USDA soil great groups at 250 *m (probabilities)* (2018).

28. Kutílek, M. & Nielsen, D. R. Soil Hydrology: Textbook for Students of Soil Science, Agriculture, Forestry, Geoecology,Hydrology, Geomorphology and Other Related Disciplines. (1994).

29. 29. Stokes, M. A. & Smiley, T. L. *An introduction to tree-ring dating*. *An introduction to tree-ring dating* (1968).

30. Holmes, R. L. Computer-assisted quality control in tree-ring dating and measurement. Tree-ring Bulletin (1983) doi:10.1016/j.ecoleng.2008.01.004.

31. Biondi, F. & Qeadan, F. A theory-driven approach to tree-ring standardization: Defining the biological trend from expected basal area increment. Tree Ring Res 64, 81–96 (2008).

32. Bunn, A. G. A dendrochronology program library in R (dplR). Dendrochronologia 26, 115–124 (2008).

33. Battipaglia, G. et al. Effects of associating Quercus robur L. and Alnus cordata Loisel. on plantation productivity and water use efficiency. For Ecol Manage 391, 106–114 (2017).

34. McCarroll, D. & Loader, N. J. Stable isotopes in tree rings. Quat Sci Rev 23, 771–801 (2004).

35. Belmecheri, S. & Lavergne, A. Compiled records of atmospheric CO2 concentrations and stable carbon isotopes to reconstruct climate and derive plant ecophysiological indices from tree rings. Dendrochronologia 63, (2020).

36. Tetens, O. Uber einige meteorologische Begriffe Z. Geophys 297–309 (1930).

37. Willmott C. J. & Feddema J. J. A More Rational Climatic Moisture Index*. The Professional Geographer 44, 84–88 (1992).

38. Allen, R. G. Evaluation of a temperature difference method for computing grass reference evapotranspiration. (1993).

39. Title, P. O. & Bemmels, J. B. ENVIREM: an expanded set of bioclimatic and topographic variables increases flexibility and improves performance of ecological niche modeling. Ecography 41, 291–307 (2018).

40. Vicente-Serrano, S. M., Beguería, S. & López-Moreno, J. I. A multiscalar drought index sensitive to global warming: The standardized precipitation evapotranspiration index. J Clim 23, 1696–1718 (2010).

41. Jevšenak, J. & Levanič, T. dendroTools: R package for studying linear and nonlinear responses between tree-rings and daily environmental data. Dendrochronologia 48, 32–39 (2018).

42. Wood, S. N. Low-rank scale-invariant tensor product smooths for generalized additive mixed models. Biometrics 62, 1025–1036 (2006).

43. Burnham, K. P. & Anderson, D. R. Model Selection and Inference: A Practical Information- Theoretic Approach. (Springer-Verlag, 2002).

44. Zabri, A. W. & Burrage, S. W. THE EFFECTS OF VAPOUR PRESSURE DEFICIT (VPD) AND ENRICHMENT WITH CO2 ON WATER RELATIONS, PHOTOSYNTHESIS, STOMATAL CONDUCTANCE AND PLANT GROWTH OF SWEET PEPPER (CAPSICUM ANNUM L.) GROWN BY NFT. Acta Horticulturae 449, 561–567 (1997).

45. Zhang, K. et al. Vegetation Greening and Climate Change Promote Multidecadal Rises of Global Land Evapotranspiration. Sci Rep 5, (2015).

46. Yuan, W. et al. Increased atmospheric vapor pressure deficit reduces global vegetation growth. Sci Adv 5, 1–13 (2019).

47. Sulman, B. N. et al. High atmospheric demand for water can limit forest carbon uptake and transpiration as severely as dry soil. Geophys Res Lett 43, 9686–9695 (2016).

48. Anderegg, W. R. L. et al. Meta-analysis reveals that hydraulic traits explain cross-species patterns of drought-induced tree mortality across the globe. Proc Natl Acad Sci U S A 113, (2016).

49. Vanoni, M., Bugmann, H., Nötzli, M. & Bigler, C. Quantifying the effects of drought on abrupt growth decreases of major tree species in Switzerland. Ecol Evol 6, 3555–3570 (2016).

50. Tonelli, E. et al. Tree-ring and remote sensing analyses uncover the role played by elevation on European beech sensitivity to late spring frost. Science of the Total Environment 857, (2023).

51. Vacchiano, G. et al. Spatial patterns and broad-scale weather cues of beech mast seeding in Europe. New Phytologist 215, 595–608 (2017).

52. Zimmermann, J., Hauck, M., Dulamsuren, C. & Leuschner, C. Climate Warming-Related Growth Decline Affects Fagus sylvatica, But Not Other Broad-Leaved Tree Species in Central European Mixed Forests. Ecosystems 18, 560–572 (2015).

53. Flexas, J. et al. Mesophyll diffusion conductance to CO2: An unappreciated central player in photosynthesis. Plant Science 193–194, 70–84 (2012).

54. Zhang, Q. et al. Response of ecosystem intrinsic water use efficiency and gross primary productivity to rising vapor pressure deficit. Environmental Research Letters 14, (2019).

55. Nock, C. A. et al. Long-term increases in intrinsic water-use efficiency do not lead to increased stem growth in a tropical monsoon forest in western Thailand. Glob Chang Biol 17, 1049–1063 (2011).

56. Peñuelas, J., Canadell, J. G. & Ogaya, R. Increased water-use efficiency during the 20th century did not translate into enhanced tree growth. Global Ecology and Biogeography 20, 597–608 (2011).

57. Mazza, G., Monteverdi, M. C., Altieri, S. & Battipaglia, G. Climate-driven growth dynamics and trend reversal of Fagus sylvatica L. and Quercus cerris L. in a low-elevation beech forest in Central Italy. Science of The Total Environment 908, 168250 (2024).

58. Li, F. et al. Global water use efficiency saturation due to increased vapor pressure deficit. Science 381, 672–677 (2023).

59. Anderegg, W. R. L., Anderegg, L. D. L., Kerr, K. L. & Trugman, A. T. Widespread drought-induced tree mortality at dry range edges indicates that climate stress exceeds species’ compensating mechanisms. Glob Chang Biol 25, 3793–3802 (2019).

60. Battipaglia, G. & Cherubini, P. Stable Isotopes in Tree Rings of Mediterranean Forests. in Stable Isotopes in Tree Rings vol. 8 605–629 (2022).

61. Medrano, H., Flexas, J. & Galmés, J. Variability in water use efficiency at the leaf level among Mediterranean plants with different growth forms. Plant Soil 317, 17–29 (2009).

62. Qi, X. et al. Growth responses to climate warming and their physiological mechanisms differ between mature and young larch trees in a boreal permafrost region. Agric For Meteorol 343, (2023).

63. Martínez-Vilalta, J. et al. Dynamics of non-structural carbohydrates in terrestrial plants: A global synthesis. Ecol Monogr 86, 495–516 (2016).

64. Merganičová, K. et al. Forest carbon allocation modelling under climate change. Tree Physiol 39, 1937–1960 (2019).

65. Lloret, F., Keeling, E. G. & Sala, A. Components of tree resilience: Effects of successive low-growth episodes in old ponderosa pine forests. Oikos 120, 1909–1920 (2011).

66. Smith, T., Traxl, D. & Boers, N. Empirical evidence for recent global shifts in vegetation resilience. Nat Clim Chang 12, 477–484 (2022).

67. Majumder, S., Tamma, K., Ramaswamy, S. & Guttal, V. Inferring critical thresholds of ecosystem transitions from spatial data. Ecology 100, (2019).

68. Fernández-Martínez, M. et al. Diagnosing destabilization risk in global land carbon sinks. Nature 615, 848–853 (2023).

69. Colangelo, M. et al. A multi-proxy assessment of dieback causes in a Mediterranean oak species. Tree Physiol 37, 617–631 (2017).

70. DeSoto, L. et al. Low growth resilience to drought is related to future mortality risk in trees. Nat Commun 11, 1–9 (2020).

71. Cabon, A., DeRose, R. J., Shaw, J. D. & Anderegg, W. R. L. Declining tree growth resilience mediates subsequent forest mortality in the US Mountain West. Glob Chang Biol 29, 4826–4841 (2023).

