## Supplementary information for "Contrasting patterns of water use efficiency and annual radial growth among European beech forests along the Italian peninsula"

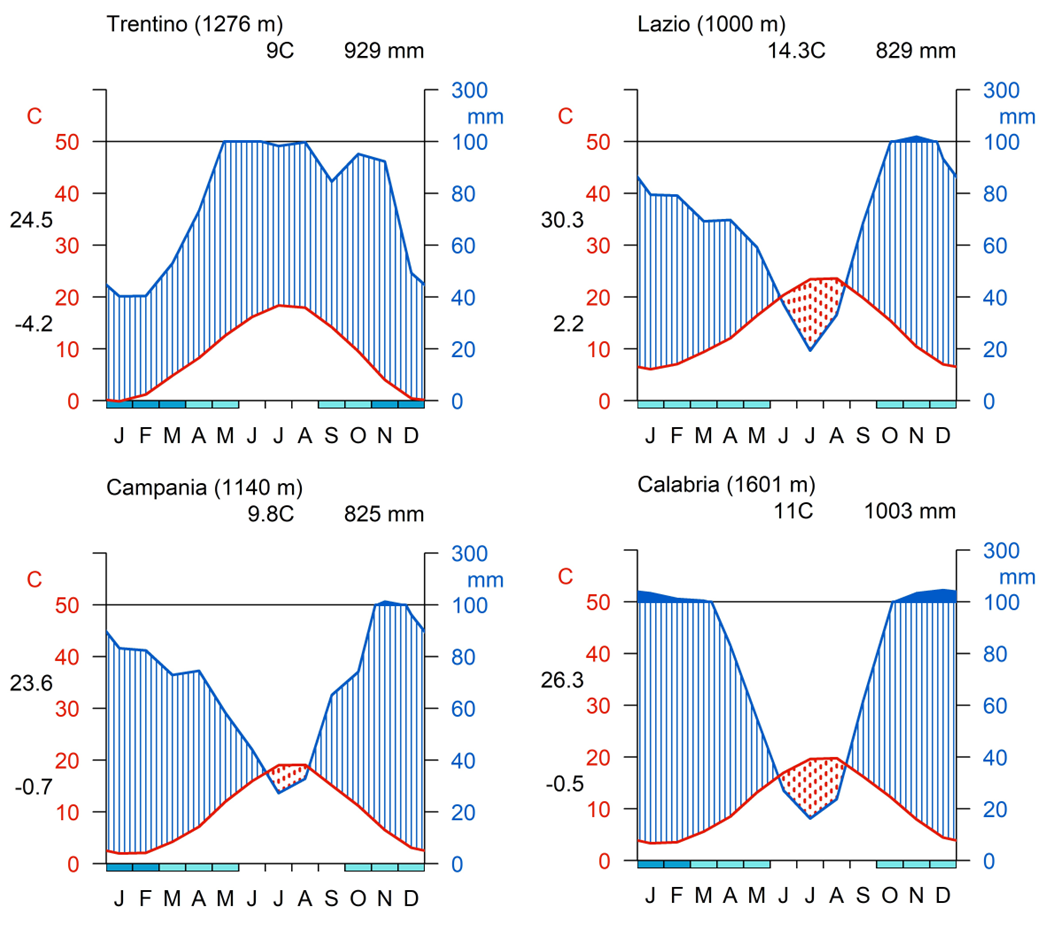

**Supplementary Fig. 1** Walter-Lieth climatograms for the four study sites: Trentino, Lazio, Campania, and Calabria (from north to south, left to right) for the period 1965-2014. The lower blue bars show the frost period.

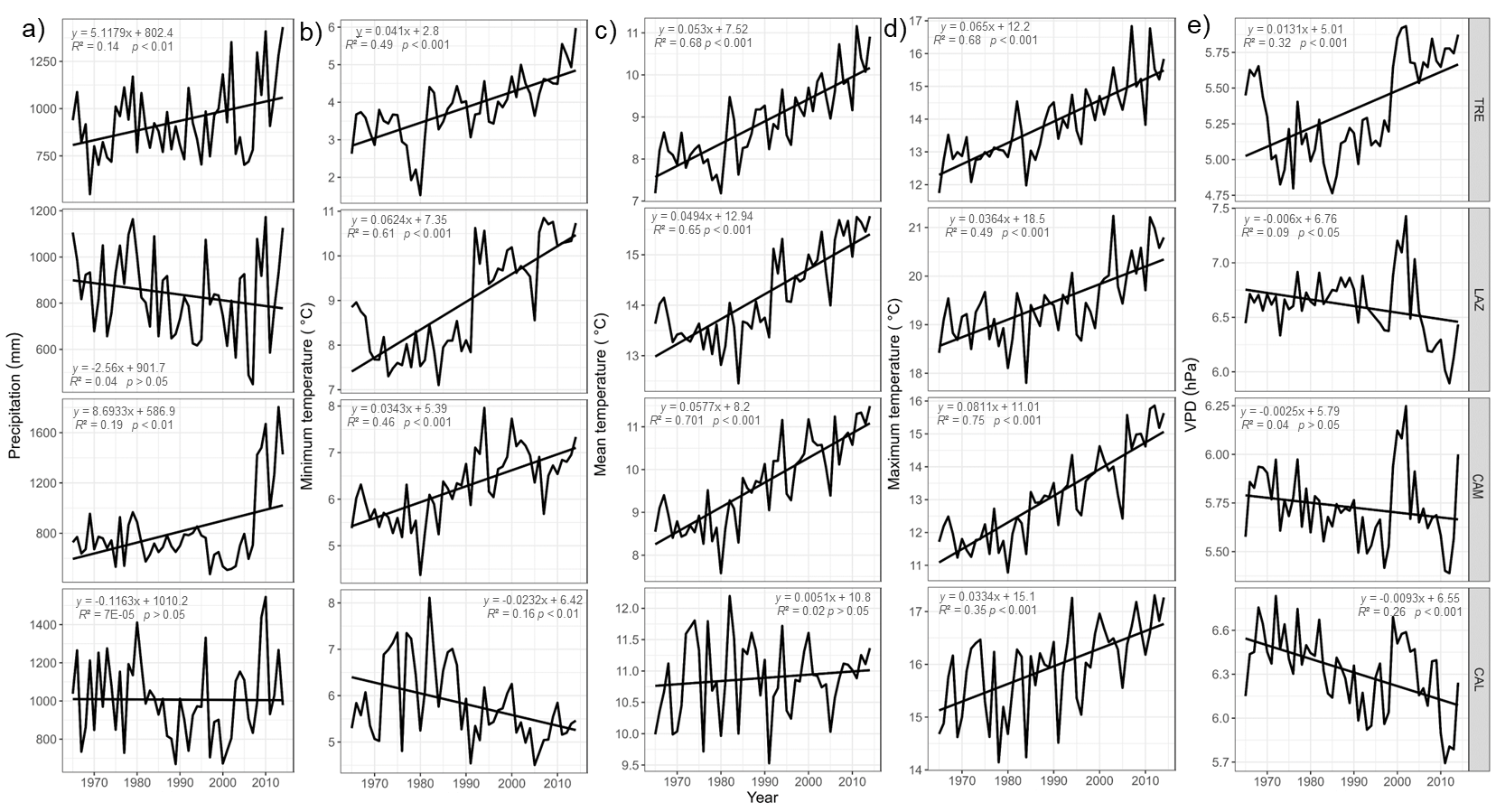
 **Supplementary Fig. 2** Climate trends at the four sites TRE, LAZ, CAM and CAL (from north to south) for the period 1965 – 2014: Mean annual a) precipitation; b) minimum temperature; c) mean temperature; d) maximum temperature and e) vapour pressure deficit (VPD). Linear regression lines are also indicated.

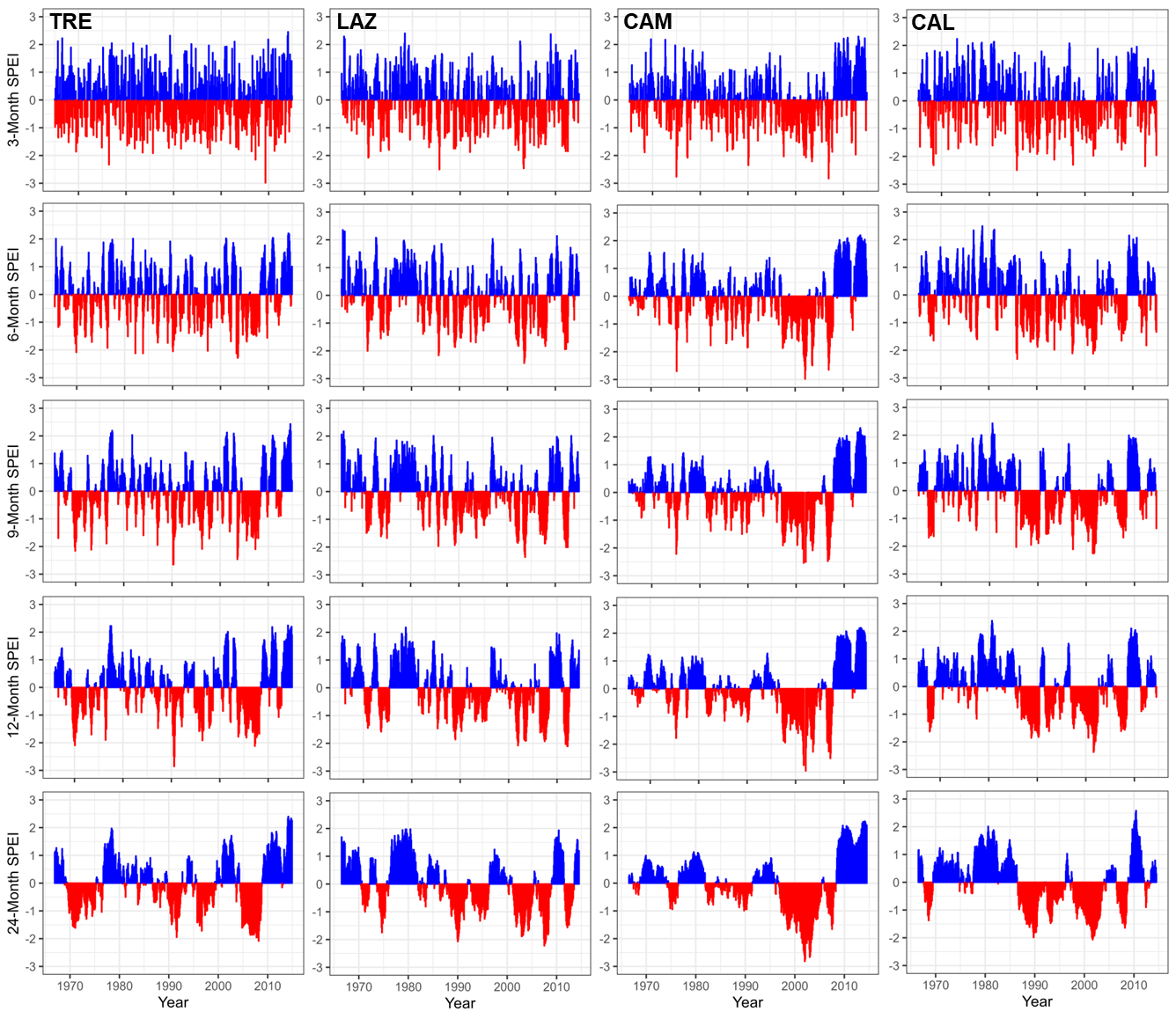

**Supplementary Fig. 3** Standardized 3-6-9-12-and 24 months SPEI at the four study sites (TRE, LAZ, CAM and CAL) for the 1965–2014 period. Negative (red) and positive (blue) values indicate drier and wetter conditions, respectively.

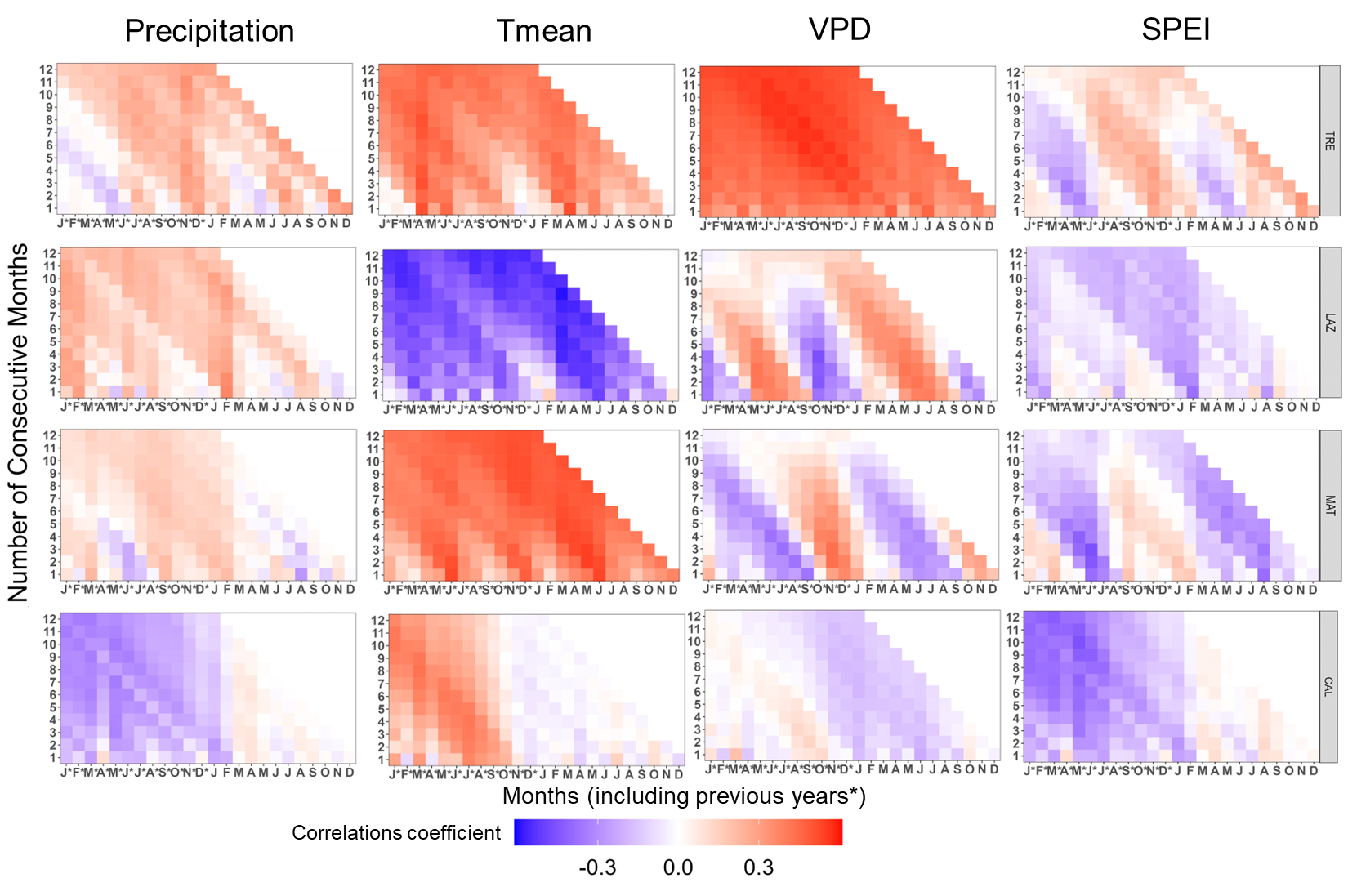

**Supplementary Fig. 4** Pearson’s running correlations between δ^13^C with monthly precipitation, mean temperature, VPD, and SPEI1 for the current and the previous year (*), over the period 1965-2014 at each site. The *y-axis* represents the time window in months. Colours (see the key) represent correlation coefficients that are significant at the level of *r* = 0.279 (*P*< 0.05).

**Supplementary Table 1** Soil type and the percentage composition of clay, silt, sand and soil water holding capacity (SWHC) at one meter depth for each site.

| Site | Soil type | Clay (%) | Silt (%) | Sand (%) | Soil water holding capacity (SWHC) |
| --- | --- | --- | --- | --- | --- |
| TRE | Udivitrands, Andisols | 19.9 | 37.8 | 42.3 | Low |
| LAZ | Haploxeralfs, Luvisols | 30.5 | 41.1 | 28.3 | High |
| CAM | Dystrudepts, Inceptisols | 30.8 | 37.8 | 31.4 | High |
| CAL | Dystrudepts, Inceptisols | 25.8 | 41.5 | 32.7 | Moderate |

**Supplementary Table 2.** Dendrochronological statistics (means ± SE) of the tree-ring width series.

| Site | No. trees | First-last years | Tree-ring width (mm) | Correlation with mean series | EPS_1_ |
| --- | --- | --- | --- | --- | --- |
| TRE | 40 | 1888-2016 | 1.01 ± 0.52 | 0.53 ± 0.11 | 1935 |
| LAZ | 20 | 1942-2014 | 2.05 ± 0.81 | 0.72 ± 0.08 | 1965 |
| CAM | 65 | 1877-2016 | 0.98 ± 0.25 | 0.51 ± 0.23 | 1910 |
| CAL | 50 | 1938-2017 | 1.43 ± 0.39 | 0.64 ± 0.09 | 1960 |

_1_The Expressed Population Signal (EPS) statistic was used to assess the strength of the common signal among the TRW series in a chronology over time. We considered an EPS > 0.85, indicating the dominance of the stand-level signal over individual tree signals. All analyses were restricted to the period covered by the youngest trees from 1965-2014.

**Supplementary Table 3** Tukey post-hoc test for TRW and age in the study sites (one-way analysis of variance with Tukey’s Honest Significant Difference test). Different letters indicate significant differences (*P* < 0.05).

| Sites | TRW (mm) | Tukey´s post-hoc | Age (years) | Tukey´s post-hoc |
| --- | --- | --- | --- | --- |
| TRE | 1.01 | *b* | 78 | *a* |
| LAZ | 2.05 | *a* | 53 | *b* |
| CAM | 0.98 | *c* | 102 | *c* |
| CAL | 1.43 | *b* | 63 | *d* |

**Supplementary TableS4** Statistic of the generalized additive mixed models for the basal area increment (BAI) trends of *Fagus sylvatica* for each site.

| Family | Model Formula | | | R^2^(adj) |
| --- | --- | --- | --- | --- |
| Gaussian | *BAI_i_=s[year_i_*(Site)]+s(age_i_)+s(SPEI18_i_)+Z_i_B_i_+ɛ* | | | 0.654 |
| Parametric coefficients |  |  |  |  |
|  | Estimate | Std. Error | t value | Pr(>\|t\|) |
| (Intercept) | 5.943 0.218 27.25 | | | <.0001 |
| Approximate significance of smooth terms: | | | | |
|  | edf | Ref.df | *F*-value | *P*-value |
| s(year):SiteTRE | 2.988 | 2.988 | 361.3 | <.0001 |
| s(year):SiteLAZ | 2.993 | 2.993 | 686.2 | <.0001 |
| s(year):SiteCAM | 2.595 | 2.595 | 233.4 | <.0001 |
| s(year):SiteCAL | 2.996 | 2.996 | 521.62 | <.0001 |
| s(Age) | 2.981 | 2.981 | 150.1 | <.0001 |
| s(spei18) | 2.839 | 2.839 | 32.0 | <.0001 |
| s(tree) | 298.39 | 303 | 79.9 | <.0001 |
